# MuClone: Somatic mutation detection and classification through probabilistic integration of clonal population structure

**DOI:** 10.1101/191759

**Authors:** Fatemeh Dorri, Sean Jewell, Alexandre Bouchard-Côté, Sohrab P. Shah

## Abstract

Accurate detection and classification of somatic single nucleotide variants (SNVs) is important in defining the clonal composition of human cancers. Existing tools are prone to miss low prevalence mutations and methods for classification of mutations into clonal groups across the whole genome are underdeveloped. Increasing interest in deciphering clonal population dynamics over multiple samples in time or anatomic space from the same patient is resulting in whole genome sequence (WGS) data from phylogenetically related samples. With the access to this data, we posited that injecting clonal structure information into the inference of mutations from multiple samples would improve mutation detection.

We developed MuClone: a novel statistical framework for simultaneous detection and classification of mutations across multiple tumour samples of a patient from whole genome or exome sequencing data. The key advance lies in incorporating prior knowledge about the cellular prevalences of clones to improve the performance of detecting mutations, particularly low prevalence mutations. We evaluated MuClone through synthetic and real data from spatially sampled ovarian cancers. Results support the hypothesis that clonal information improves sensitivity in detecting somatic mutations without compromising specificity. In addition, MuClone classifies mutations across whole genomes of multiple samples into biologically meaningful groups, providing additional phylogenetic insights and enhancing the study of WGS-derived clonal dynamics.

## Introduction

Genomic accumulation of somatic point mutations, or single nucleotide variants (SNVs) can disrupt the regular activity of cells and may result in cancer initiation and progression. Collectively, the complete repertoire of SNVs across a cancer genome (numbering in the thousands) form a statistically robust marker for inferring clonal populations and studying tumour evolution. As such, accurate detection of all somatic SNVs, including those with low prevalence, is vital as they can define clones with phenotypic properties of interest. Mechanistic association of specific clones with properties such as treatment resistance, metastatic potential and fitness under therapeutic selective pressures remains a key objective of biomedical investigators studying tumour progression.

Phylogenetic analysis can encode the evolutionary lineage of tumour cells across time and anatomic space [7, 10, 14, 16, 17, 23, 25]. [6] sequenced multiple spatially separated samples from renal cell carcinomas and related metastatic sites to reveal the evolutionary patterns. Samples were related through phylogenetic analysis and distinguished at a coarse level mutations that were shared and ancestral from those that occurred in subsets of cells. In a subsequent lung cancer study, 25 regions from seven non-small sections of malignant patients were sequenced [2] and more recently the TRACERx study involving 100 lung cancer patients with 3 samples per patient has reported genomic instability as a determinant of treatment response [10]. Our recent work has determined clonal population dynamics over time in breast cancer xenografts [5], follicular lymphoma timeseries sampling across clinical trajectories [14] and anatomic space in intraperitoneal sites of primary high grade serous ovarian cancer [16], showing that the relative composition of constituent clones in multi-sample studies provides major insight into disease spread.

In the limit case, all cells likely harbour unique genomes, however due to the nature of branched evolutionary processes, clones can be coarsely modeled as major clades in the cell lineage phylogeny of a cancer. These clades share the majority of mutations, and therefore define first approximations to the genotypes of clones. Clonal genotypes and their relative abundances in the cancer cell population can be approximated by clustering mutations measured in bulk tissues and estimating the cellular prevalences (the variant fraction of tumour cells) [20, 26].

Phylogenetic algorithms mostly use mutations (represented as binary genetic markers), as inputs to infer the branched evolutionary lineages of tumour cells [3, 18]. Thus, accuracy of mutation detection will impact the performance of phylogenetic inference algorithms.

Detection of low prevalence mutations is a major challenge due to weak signal to noise ratio, owing to: (i) impure samples which are contaminated by normal cells, (ii) copy number alteration of the genome, and (iii) the presence of mutations in only a fraction of tumour cells (intra-tumour heterogeneity). We assert in this work that prior knowledge of clonal population structure will improve detection of mutations defining low prevalence clonal genotypes.

### 0.1 Previous work

SNV calling algorithms are ubiquitous in the literature, but the problem remains challenging particularly for detecting low prevalence mutations. Algorithms have been developed for calling mutations from a single sample [8, 13], paired (matched normal and tumour) samples [1, 4, 12, 19, 22], or multiple samples [11, 24]. [4] uses a feature based classifier called Mutationseq for calling mutations. The features are constructed from matched paired normal and tumour samples. [22] introduced Strelka a method for somatic SNV and small indel detection from sequencing data of matched normal and tumour samples. It is based on a Bayesian approach which uses allele frequencies for both normal and tumour samples with the expected genotype structure of the normal. [1] proposed Mutect which uses a Bayesian classifier to detect mutations from matched normal and tumour samples. It uses various filters to ensure high specificity. [11] proposed multiSNV which jointly considers all available samples under a Bayesian framework to improve the performance of calling shared mutations. [21] and [24] refine and correct the SNV calls from GATK [15] using the phylogeny information across multiple samples.

### 0.2 Our contribution

In MuClone, we exploit prior knowledge of tumour clone prevalence information and copy number inference across multiple samples to improve the performance of detecting mutations, with the goal of better detecting low prevalence clones. In this contribution, the clonal information is provided by running PyClone on the data [16, 20], and copy number information is estimated by TITAN [9]. However we note that the model can be applied to clonal and copy number data obtained from any method. In addition, Muclone classifies mutations into clusters sharing similar cellular prevalence which provides the opportunity of profiling their dynamic across time or space and adds a rich layer of interpretation into the detection process.

We tested MuClone through simulation studies and an application to real, multiple sample, patient data. These experiments reveal that incorporating the cellular prevalences of different clones improves accuracy. Moreover, in real data MuClone exhibits higher sensitivity in detecting mutations without compromising specificity compared with other methods.

## 1 Notation

We begin by introducing notation used in the MuClone model. For each locus *n* = 1,*…,N*, the samples are indexed from *m* = 1*…M*.

We assume that bulk tumour DNA arises from three populations: (i) normal cells; (ii) malignant cells without a mutation of interest (reference); and (iii) malignant cells with the mutation of interest (variant). The genotype for locus *n* is denoted 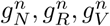, for normal, reference, and variant populations, respectively.

The symbol *A* denotes the allele that matches the reference genome; conversely, the symbol *B* denotes the allele that does not match. Since human genomes usually have two copies of DNA content, the genotype of a diploid locus is one of *AA*, *AB* or *BB*. If there is a coincident copy number alteration, the possible genotypes change accordingly. Each of the genotype variables 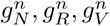 take values in 𝒢 = {−,*A,B,AA,AB,BB,AAA,AAB,…*}, for example, the genotype *AAB* refers to a genotype with two reference alleles and one variant allele. At each locus *n* of sample *m*, the genotype state is represented as the ordered list 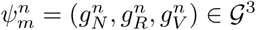, which represents the genotype in each population.

For example, *c*(*AAB*) = 3 and *b*(*AAB*) = 1. The symbol − denotes the genotype with no alleles, in other words, a homozygous deletion of the locus.

In sample *m*, 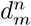 is the total number of reads aligned (read depth) at locus *n* and 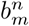 is the number of aligned reads with *B* alleles. The tumour content, defined as the fraction of cancer cells in sample *m*, is denoted by *t*_*m*_. We subdivide cancerous cells into cells from the variant population and those that are from the reference population. The proportion of cells that are from variant population is called the cellular prevalence.

Previously known clonal information from, for example, PyClone [16, 20] is encoded in an ordered list π’= (ϕ’,*τ*’). The cellular prevalence is recorded in the *M* × *K* matrix ϕ’, where the (*m,z*) element, 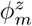, represents the cellular prevalence of the *m*th sample and *z*th clone. The *K*-vector *τ*’represents prior the clonal prevalence. In PyClone’s case, the clonal prevalence of the *z*th clone is the empirical proportion of the number of mutations in the *z*th clone to the total number of mutations clustered. Since our interest lies in calling mutations, and many statistical models for inference of clonal population structure, including PyClone, only consider somatic mutations, we extend π’to include a wildtype clone *z* = 0 and denote the resulting list as π = (ϕ,*τ*). In particular, we add a column of wildtype cellular prevalences to ϕ’to create ϕ, and add a wildtype clone prevalence to the vector *τ*’; *τ* is formed by normalizing (*τ*_0_,*τ*’).

## 2 MuClone

MuClone uses previously known cellular prevalence information to improve mutation detection and classification. For each sample, MuClone detects mutations from joint analysis of multiple samples. We encode this process in a generative probabilistic framework to perform joint statistical inference of multiple observations (from multiple samples) of the variant allele counts of a mutation of interest.

The probabilistic graphical model of MuClone is depicted in Figure 1.

**Figure 1.**
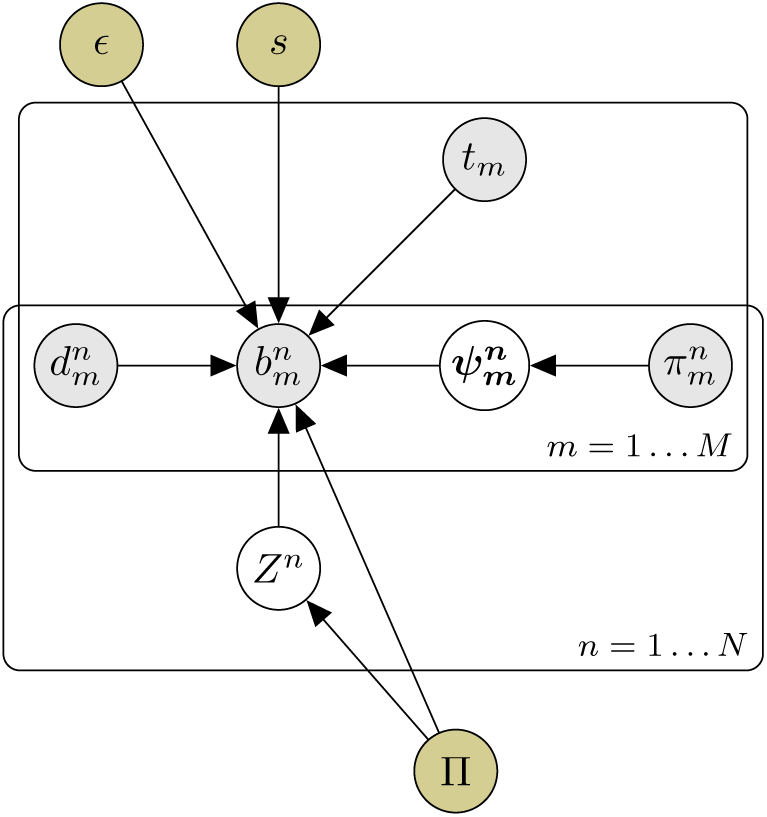
Probabilistic graphical model of MuClone: white nodes are unobserved variables; grey shaded nodes are observed variables; golden nodes are external information. The variables *m* ∈{1*…M*} and *n* ∈{1*…N*} index the samples and the loci respectively. In sample *m*, 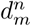 is the total number of reads aligned at locus *n* and 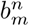 is the number of aligned reads with *B* alleles. The genotype state is 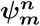 and 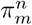 is the prior over the genotype states. The tumour content of sample *m* is *t*_*m*_ and the error rate is. The parameter *s* stands for the precision parameter. The clonal information is π and the variable *Z*^*n*^ denotes the mutation clone.

### 2.1 Model definition

For simplicity, we first assume that the number of reads containing the variant alleles at a given locus follows a Binomial distribution with genotype specific variant probability *p*(*g*) and read depth 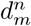

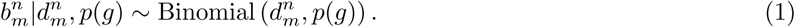

The variant probability *p*(*g*) : 𝒢 → [0,1] is defined as

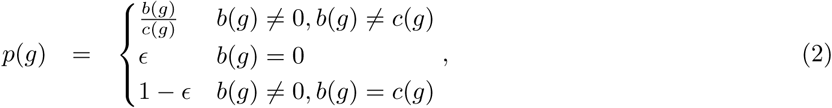

where *g* is the genotype and ε > 0 is a small positive constant that accounts for sequencing error. It allows for non-zero variant reads, due to sequencing error, when there are no variant alleles in genotype *g*.

However, since the sequenced reads are independently sampled from an infinite pool of DNA fragments, each read may belong to the normal, reference, or variant population. Therefore, using a single genotype state, *g*, introduces error into our analysis. To account for this fact, we consider using the full genotype state, 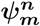, at a given locus *n* to model the number of variant reads.

The variant allele probability for the *n*th locus in the *m*th sample from the *z*th clone, denoted by 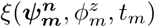, is proportional to the sum of the (properly scaled) variant probabilities from each population:

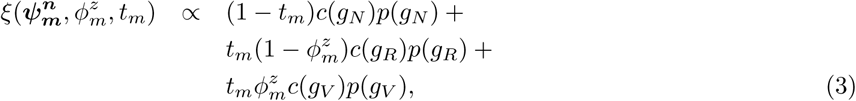

where the first term (1 − *t*_*m*_)*c*(*g*_*N*_)*p*(*g*_*N*_) is proportional to the probability of sampling a read containing variant allele from the normal population, and the second and third terms, 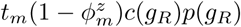 and 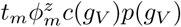, are proportional to the probabilities of sampling a read containing variant alleles from the reference and variant populations, respectively.

Considering the full genotype state, the number of reads containing the variant alleles at a given locus *n* that belongs to clone *Z*^*n*^ follows a Binomial distribution with probability

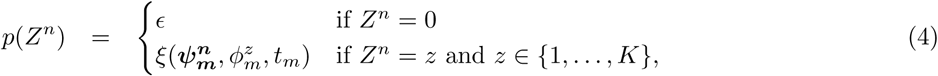

where ∊ accounts for sequencing error in wildtype clone and 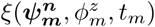 is the variant alleles probability for *n*th locus, *m*th sample from *z*th clone. According to Equation (3), tumour content and cellular prevalence information which is encoded in π, are incorporated to estimate 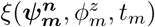.

Since empirical evidence shows that variant read data is overdispersed, we replace the Binomial model (1) with a BetaBinomial model

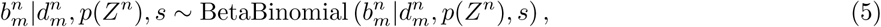

where *p*(*Z*^*n*^) is the expected variant alleles probability and the hyperparameter *s* is the precision parameter of the BetaBinomial distribution. The BetaBinomial distribution in Equation (5) assigns a small chance for mutation when the locus is wildtype, otherwise it is governed by the prior clonal information.

To fully express our model, for each locus, we assume the genotype state follows a categorical distribution with probability vector 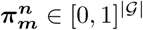 whose *i*th element is the probability of the *i*th genotype state

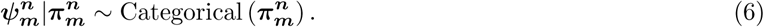

The number of possible genotype states, denoted by |𝒢|, is finite given the copy number information. For simplicity, we assume every element of 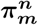 is equal to 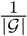.

In addition, we also assume that the clonal assignment of a locus, denoted by *Z*^*n*^, follows categorical distribution with probability vector *τ*:

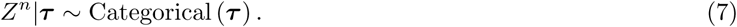

Our probabilistic framework can be succinctly written as

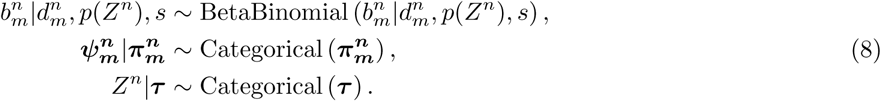

### 2.2 Inference

Based on the generative model introduced in (8) mutations are inferred via the posterior probability distribution of a locus *n* belonging to clone *z*, 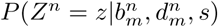 is:

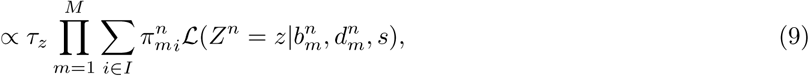

where the variable *i* indexes 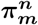 over the genotype states, *I* = {1*…*| 𝒢|}. The posterior probability of locus *n* belongs to clone *z* is proportional to the likelihood of observing 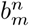 number of nucleotides matching the variant alleles times the prior over tumour clone *z*. The likelihood function, 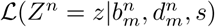, is the BetaBinomial distribution defined in (5).

Based on basic decision theory, a decision can be extracted from a posterior distribution given a loss function. Under the loss function *l*(*z,z*’) = **1**[**1**[*z* = 0] ≠ **1**[*z*’= 0]], the decision is simply the maximum a posteriori (MAP). That is, if the probability *η* of belonging to any of the tumour clones is greater than 0.5, we conclude that the locus is mutated in at least one of the *M* samples. The value of *η* is

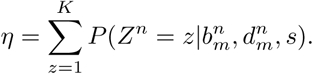

If locus *n* is mutated in at least one of the *M* samples, then the probability of mutation in each sample is calculated separately as

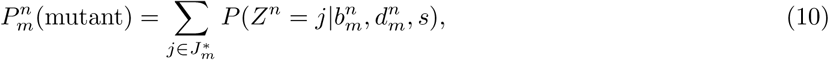

Where 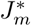 is the set of clones of sample *m* whose cellular prevalences are greater than a fixed positive threshold called ϕ_*T*_,

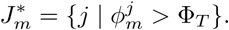

The threshold ϕ_*T*_ distinguishes the clones of sample *m* in which their non-zero cellular prevalence are due to actual variant alleles. The default value of ϕ_*T*_ is zero. However, depending on the method used for estimating cellular prevalences, it can be set to another positive value, if some non-zero input cellular prevalences indicate wildtype clones.

In addition, MuClone assigns the locus to clone *z*\* that maximizes

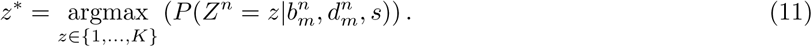

This classifies mutations to one of the previously known clones. The classification of mutations helps in biological interpretation and phylogenetic analysis of the data.

## 3 Experimental Result

We evaluated MuClone on both synthetic and real data.

### 3.1 Synthetic data

#### 3.1.1 Data generation

Synthetic data was generated for *N* number of loci, and *M* samples. We varied the number of tumour clones, *K*, sequencing error rate, ∊, and tumour content, *t*_*m*_, for each sample.

The cellular prevalences of tumour clones were sampled from a Uniform distribution over the closed interval [0,1] such that all clones are not present in all samples. Loci were randomly assigned to different clones. Then, for each locus in each sample, the coverage was sampled from a Poisson distribution with the mean *d*_*m*_. Wildtype copy number was deterministically set to 2 and a copy number profile (major and minor copy number) was generated by the following steps: First, total copy number, *C*^*t*^, was sampled from integers between 1 and *C*_max_. Second, an integer number, *C*^*b*^, was randomly picked from 1 to *C*^*t*^ and *C*^*a*^ was defined as *C*^*a*^ = *C*^*t*^ − *C*^*b*^. Last, major copy number was set to the maximum of *C*^*b*^ and *C*^*a*^; minor copy number was set to the minimum of those two values. Then, corresponding to each clone, the number of variant reads were sampled from the Beta-Binomial distribution described in Equation (5).

#### 3.1.2 Synthetic data evaluation

We simulated synthetic data for 200 loci from 4 samples of a patient, with 10 underlying clones, including an ancestral clone. The maximum copy number was 5, and error rate was 0.01. The average depth of sequencing was assumed to be 100. This process was repeated 10 times in order to compute distributions over accuracy metrics. The distribution over clone prevalences in multiple samples across different runs, and the cellular prevalence of samples in a random experiment are depicted in Figures S1 and S2 respectively. As shown, similar to real data scenario, not all of the clones are present in all of the samples.

We began evaluation by investigating the effect of systematically ‘shielding’ MuClone from clonal information (Figure 2). Clonal information was perturbed by (i) adding noise to the cellular prevalences of tumour clones, or (ii) removing clonal information. The noise was generated from a normal distribution with different standard deviations (sd): 0, 0.01, 0.1, and 0.25. The noise value was randomly either added to or subtracted from the cellular prevalence of the clone, while bounding the resulting value between 0 and 1.

**Figure 2.**
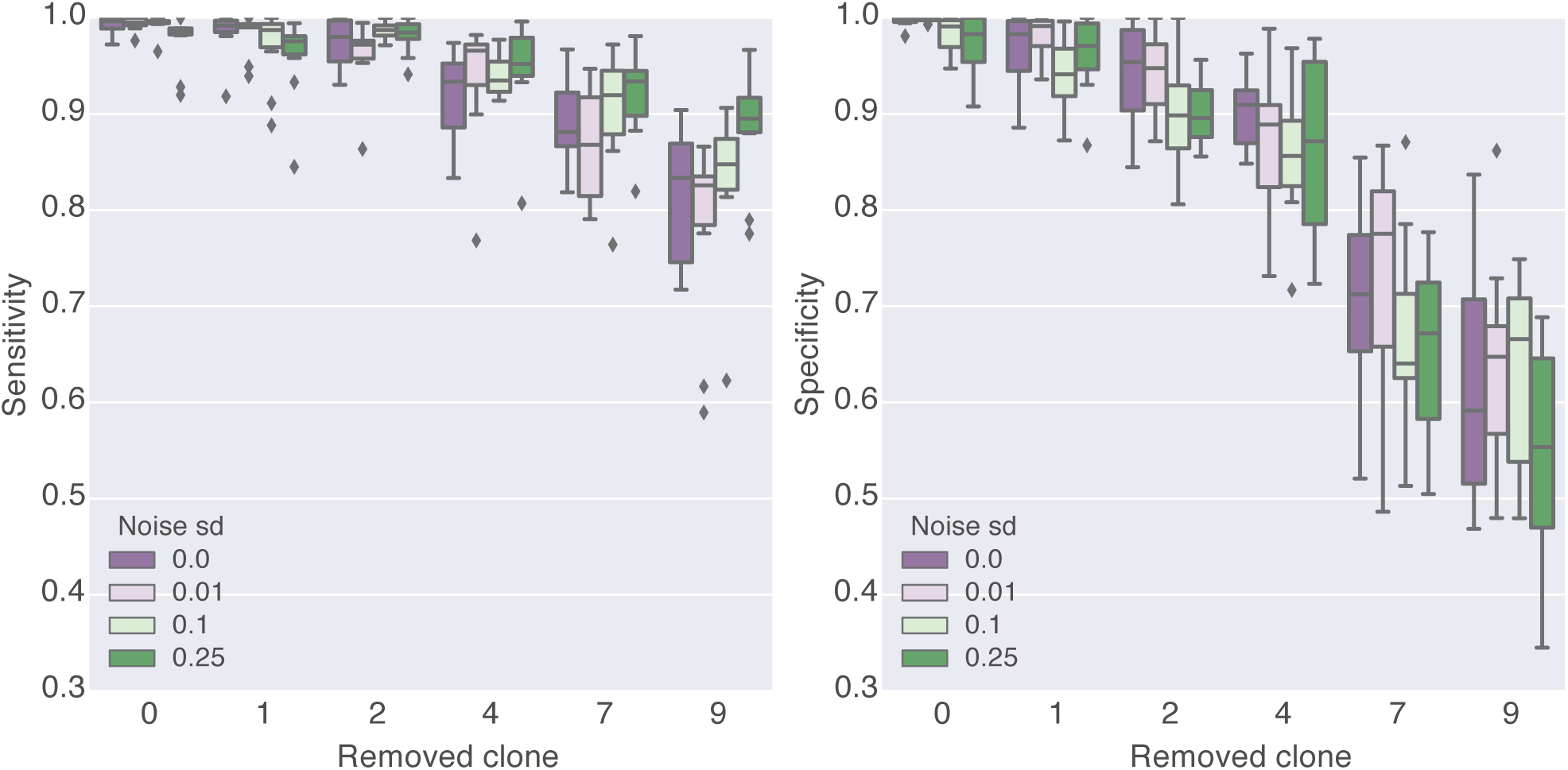
MuClone’s sensitivity and specificity with inaccurate clonal information. The synthetic data is generated for 200 loci from 4 samples of a patient, with 10 underlying clones. Maximum copy number is 5, and error rate equals 0.01. The average depth of sequencing is about 100. Noise sd is the standard deviation of noise added to (or subtracted from) the clones cellular prevalences. Removed clone is the number of clones that their clonal information is not accessible to MuClone. MuClone’s parameters: Wildtype prior is 0.5, ϕ_*T*_ is 0.02, error rate is 0.01, tumour content is 0.75, and precision parameter equals 1000.

As expected, both sensitivity and specificity were highest when clonal information was most complete and most accurate (Figure 2). This suggests that clonal information can indeed improve the accuracy of detecting mutations and establishes the theory of MuClone’s approach. Furthermore, sensitivity and specificity were only marginally impacted by the added noise in the clonal information, suggesting MuClone should be able to cope with the modest levels of erroneous information in the prior.

Naturally, accuracy was most severely impacted when the least amount of clonal information was input (Figure 2). For different sd values, the sensitivity and specificity of removing various number of clones were compared through Kruskal-Wallis test (p values ≤ 4e-5 and ≥ 1e-9) which shows the change in performance is significant when perturbing the quantity and quality of clonal information.

We next explored how sensitivity and specificity changes as a function of the wildtype prior and ϕ_*T*_ values (Figure 3). We generated data by setting the wildtype prior at 0.5, 0.75, and 0.99; and ϕ_*T*_ at 0.001, 0.01, 0.02, 0.03, 0.04, and 0.05. MuClone’s sensitivity and specificity were near 1 for ϕ_*T*_ about 0.02. Sensitivity dropped with higher ϕ_*T*_ because mutations were miscalled as wildtypes and specificity dropped with lower ϕ_*T*_ because wildtypes were miscalled as mutations. The optimal ϕ_*T*_ was about 0.02 when wildtype prior equals 0.5. These parameters were used in the following experimental results.

**Figure 3.**
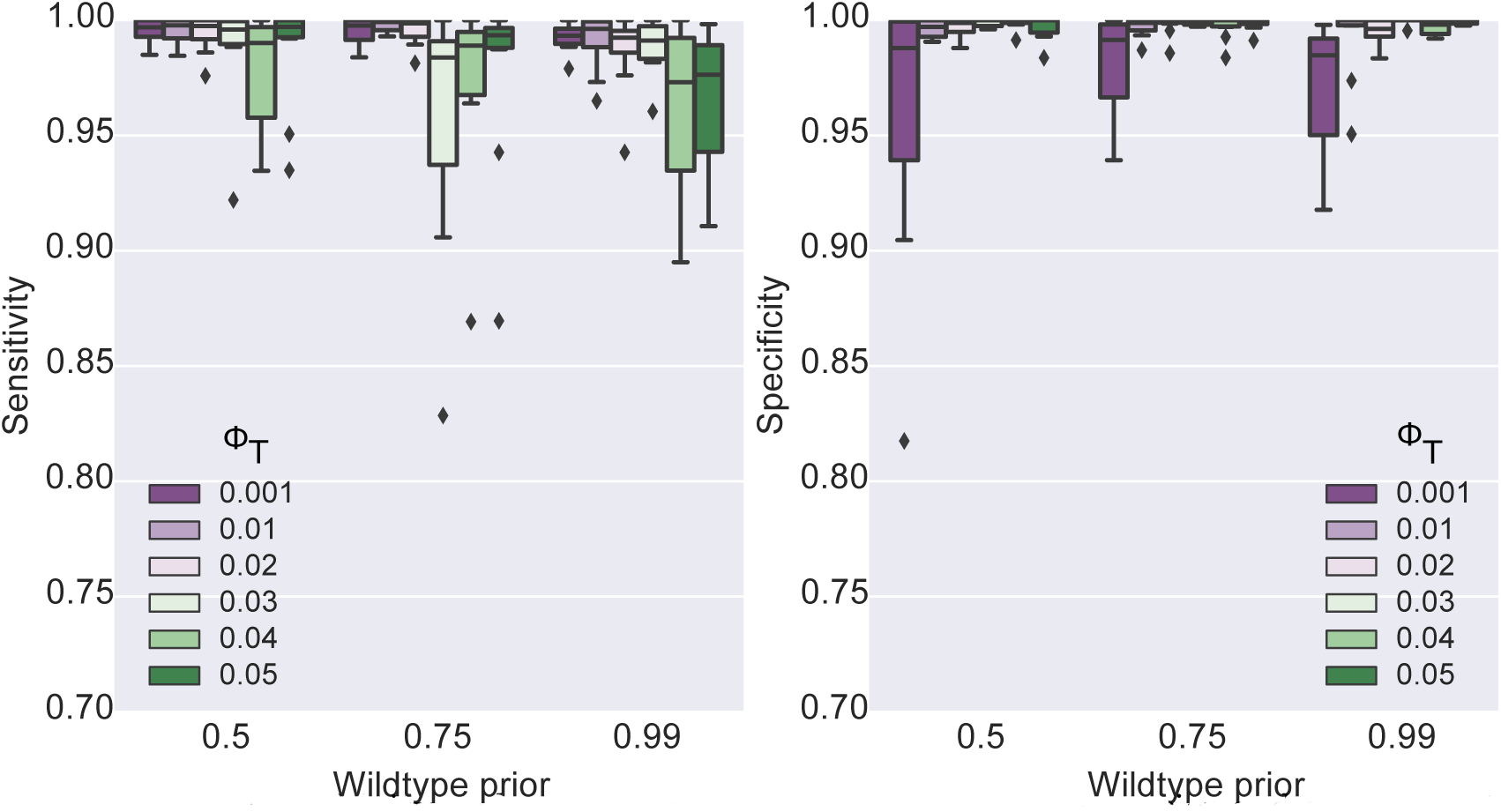
MuClone’s sensitivity and specificity at three different wildtype prior and six different ϕ_*T*_ values. The synthetic data is generated for 200 loci from 4 samples of a patient, with 10 underlying clones. Maximum copy number is 5, and error rate equals 0.01. MuClone’s parameters: error rate is 0.01, tumour content is 0.75, and precision parameter equals 1000.

The performance of MuClone was tested with various tumour content (from 0.1 to 0.99) and different error rates (0.01 and 0.001) (Figure 4). For samples with tumour content greater than 0.5, sensitivity remained slightly less than 1 and specificity near 1. Sensitivity and specificity dropped to only about 0.9 when the tumour content in the sample was as low as 0.1, establishing promising performance over different ranges of tumour content with different error rates (likely scenarios in real data).

**Figure 4.**
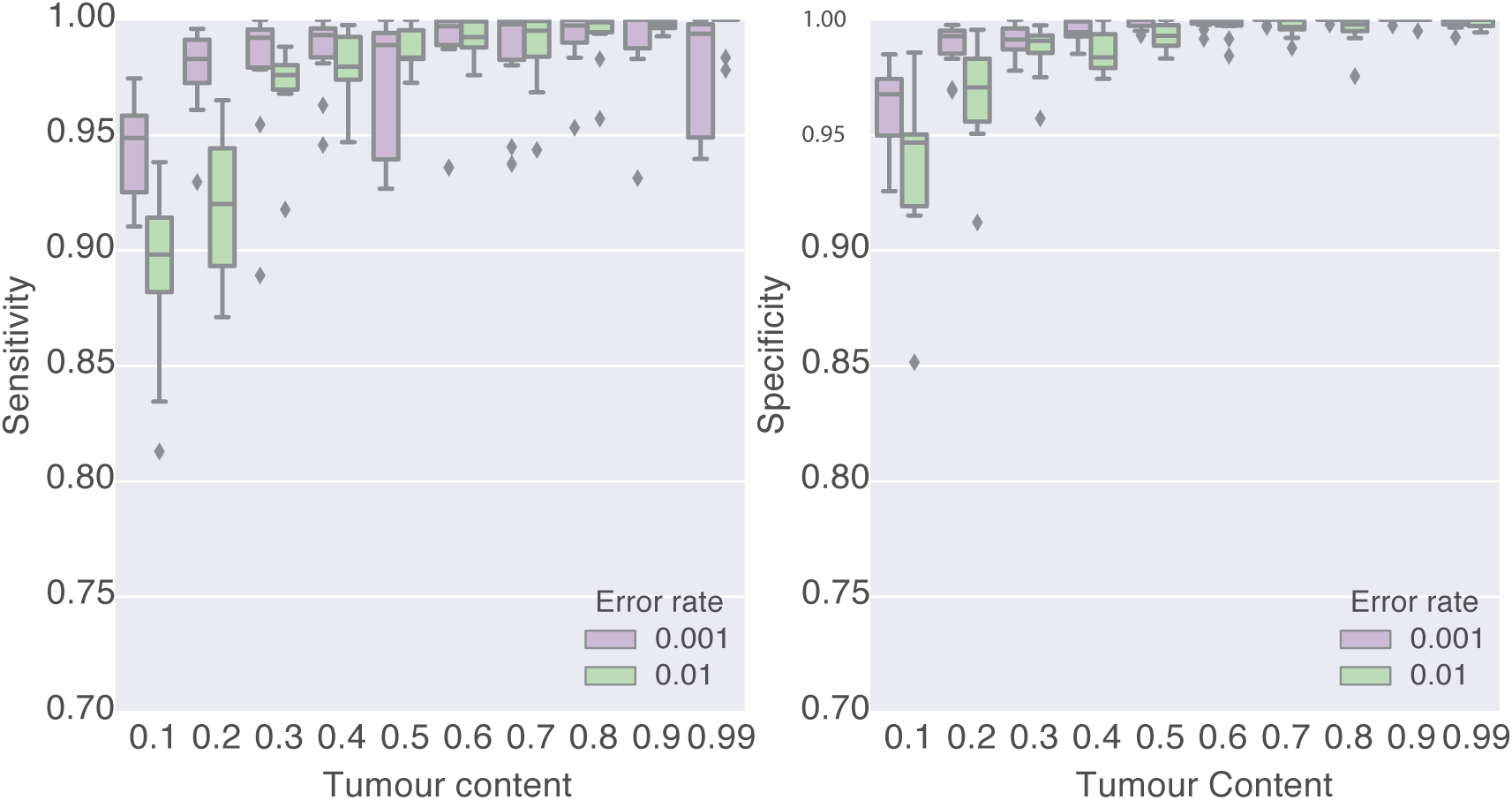
MuClone’s sensitivity and specificity with two different error rate values and tumour content values from 0.1 to 0.99. The synthetic data is generated for 200 loci from 4 samples of a patient, with 10 underlying clones. Maximum copy number is 5. MuClone’s parameters: Wildtype prior is 0.5, ϕ_*T*_ is 0.02, and precision parameter equals 1000.

Figure 5 demonstrates how well the mutations were classified by MuClone. The input clonal information had been perturbed by adding noise with standard deviation of 0.01 to simulate a more realistic scenario. In Figure 5(a), each bin (*i,j*) shows the number of mutations belonging to clone *i* that MuClone classified them into clone *j*, divided by the total number of mutations. Figure 5(a) shows 89% of mutations were classified into the right clone as the diagonal elements are larger than the other ones.

**Figure 5.**
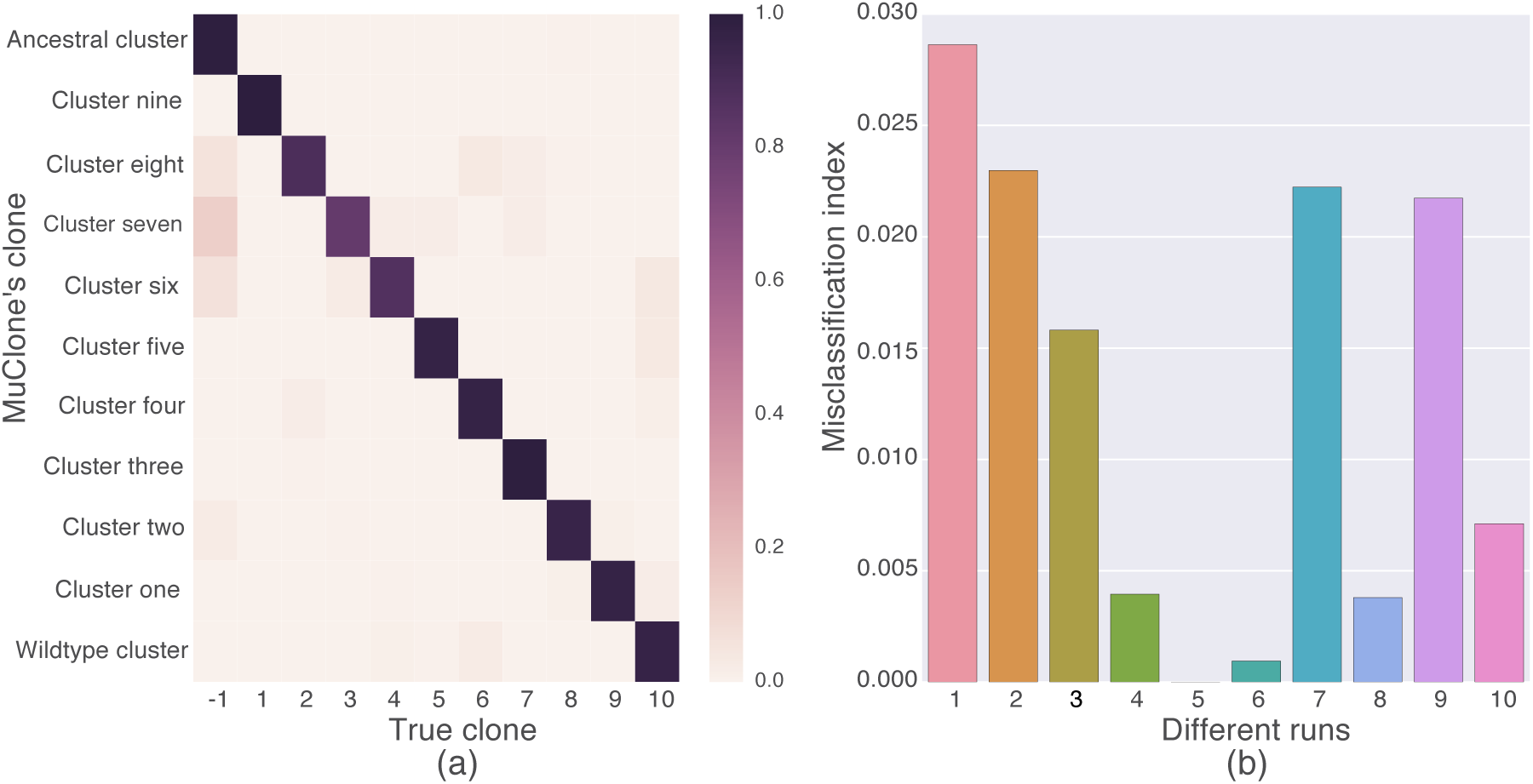
Evaluate the classification of mutations in different clones by MuClone. The synthetic data is generated for 200 loci from 4 samples of a patient, with 10 underlying clones. Maximum copy number is 5, error rate is 0.01, and noise sd is 0.01. MuClone’s parameters: Wildtype prior is 0.5, ϕ_*T*_ is 0.02, error rate is 0.01, and precision parameter equals 1000. (a) Bin (*i,j*) shows the normalized number of mutations in clone *i* but classified in clone *j*. Diagonal elements show 89% of the mutations are classified correctly. (b) Misclassification index for 10 independent experiment runs.

In order to show that the classification errors have occurred between clones with small phylogenetic distance, we define misclassification index calculated as below:

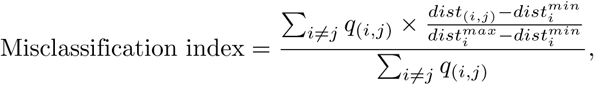

where *q*_(*i.j*)_ is the number of mutations in clone *i* that have been classified into clone *j*. The Euclidean distance between the cellular prevalences of clone *i* and *j* is *dist*_(*i,j*)_. The distance of the closest and farthest clone to clone *i* is denoted by 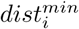 and 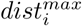 respectively. In Figure 5(b), small misclassification indexes demonstrate that misclassified mutations occur between close clones which can be interpreted as recently separated clones considering their

### 3.2 Real data

We next tested MuClone’s performance on whole genome sequencing data (30*X*) from multiple tumour samples surgically resected from high grade serous ovarian cancer patients [16]. The samples were obtained from different spatially distributed metastatic sites. Brief details about the number of samples for each patient, sample sites and the number of validated loci for each patient are shown in Table S1.

The clonal information and experimentally re-validated mutations status were taken from [16], estimated from running PyClone on the deep targeted sequencing data (*>* 1000x coverage) from the same samples and in three patients with accompanying single cell sequencing data (see Table S16 in [16]). In order to eliminate germlines, loci with any number of variant nucleotides in the corresponding normal sample were removed from the data set. Then, the performance of MuClone was benchmarked against Strelka, MutationSeq, Mutect, MultiSNV and naive MuClone. Naive MuClone is a version of MuClone where no clonal information is provided (which assumes that all mutations are from the ancestral clone).

The performance of MuClone is compared with other methods in Figure 6. The Youden’s index, sensitivity and specificity was averaged across different samples of 7 patients. And each patient’s performance is shown separately in Figures S3 to S9. Youden’s index is calculated as:

**Figure 6.**
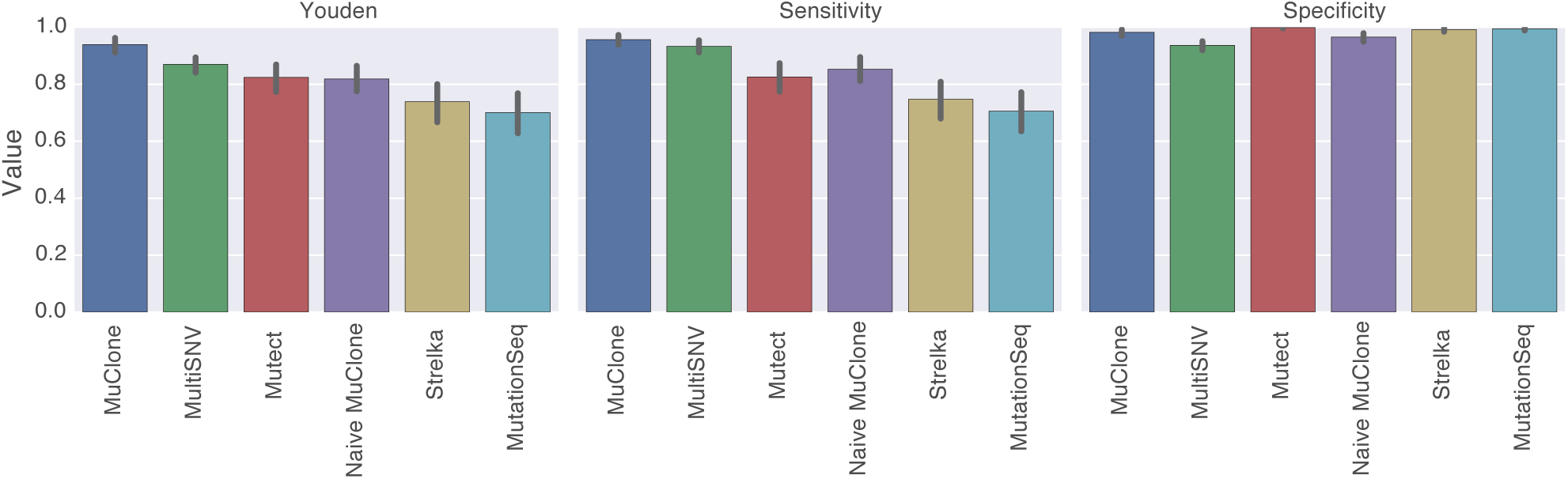
The comparison of the Youden’s index, sensitivity and specificity of different mutation detection methods. MuClone’s parameters: Wildtype prior is 0.5, ϕ_*T*_ is 0.02, error rate is 0.01, and precision parameter equals 1000.

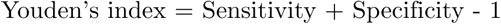

In aggregate, MuClone outperforms other methods by improving sensitivity without compromising specificity (Figure 6). For each patient, the receiver operating characteristic (ROC) curves are shown in Figures S10 to S16. The detailed number of miscalled loci are listed in Table S2. False negatives are mainly because the WGS data is under-represented (the average depth of the WGS data is about 30X) and lacks any variant alleles that are present in the targeted sequencing data. The false positives are mostly because of technical artefacts.

In Figure 6, Strelka, MutationSeq, Mutect and Naive MuClone have lower performance as they do not incorporate the information across multiple samples for calling mutations. We calculated Welch’s t-test for unequal population variances on MuClone and MultiSNV Youden’s index results. The performance of MuClone was statistically higher than MultiSNV (p-value = 0.0006). Importantly, MuClone improves sensitivity, enabling the detection of more mutations across whole genome. Figure 7 depicts the classification of mutations into clones relative to ground truth, as defined by running PyClone on the data (omitting singleton clusters [16]). Each bin shows the normalized number of mutations. 93% of the elements in Figure 7 are diagonal which means MuClone classifies them correctly. Misclassification index for patient 1 is 0.015 which implies that misclassified mutations are classified into phylogenetically close clones.

**Figure 7.**
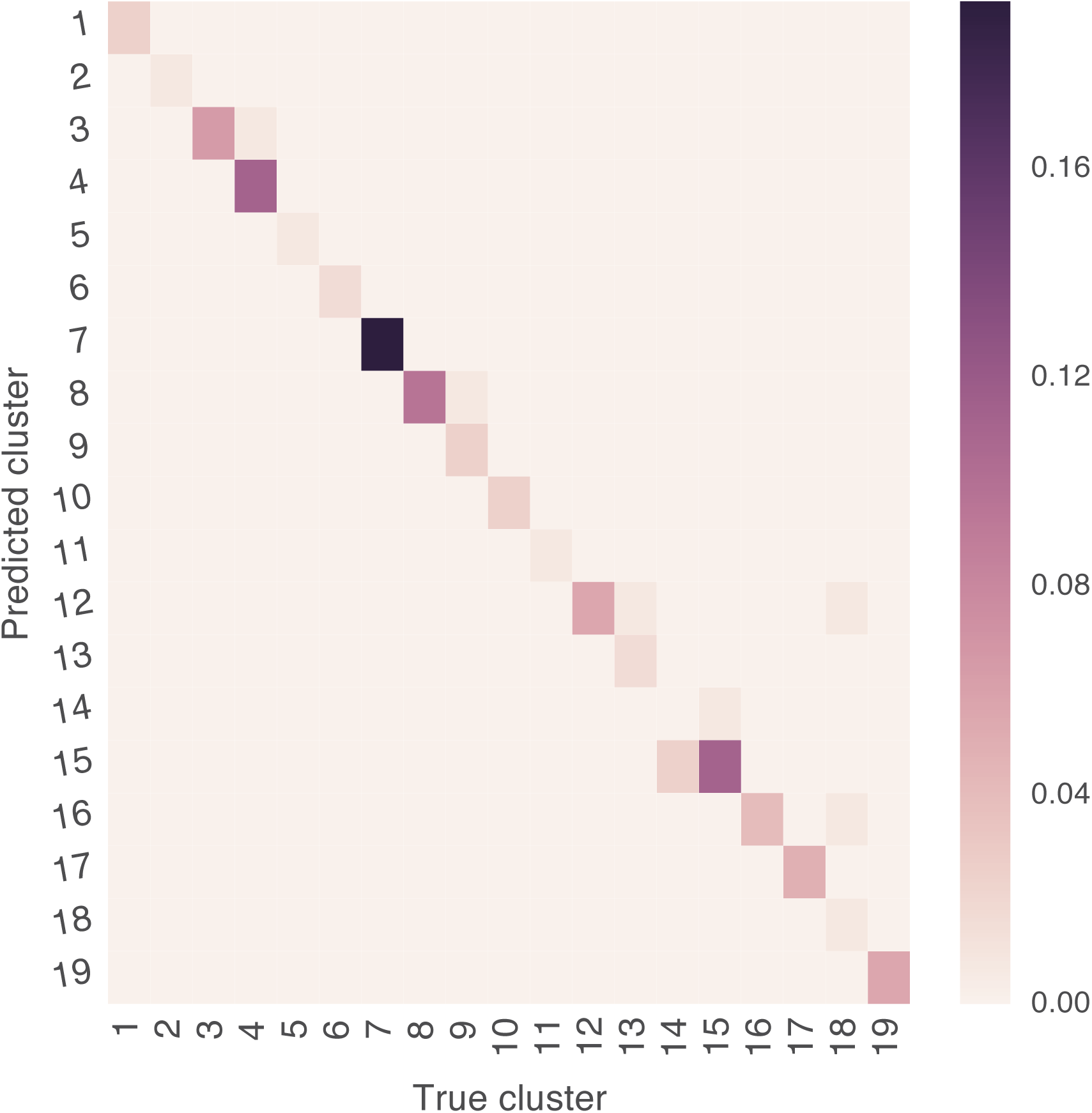
Classification of 153 mutations of patient 1 across 6 samples. Bin (*i,j*) shows the number of mutations in clone *i* that were classified into class *j* by MuClone, divided by the total number of mutations. MuClone’s parameters: Wildtype prior is 0.5, ϕ_*T*_ is 0.02, error rate is 0.01, and precision parameter equals 1000. 93% of the elements are diagonal.

## 4 Discussion and Conclusion

We studied the use of clonal information for the purpose of somatic mutation detection and classification in multi-sample whole genome sequencing data. The proposed statistical framework uses the clones cellular prevalences and copy number information for detection and classification of low prevalence mutations. MuClone outperformed other popular mutation detection tools while providing the added benefit of classifying whole genome sequencing mutations into biologically relevant groups. Both simulation and real data results showed the cellular prevalences of tumour clones are beneficial information for improving the sensitivity. Importantly, our results suggest improvement in sensitivity can be achieved without compromising specificity. As accuracy of detecting mutations can affect the performance of phylogenetic analysis, we suggest this improvement will impact the field of multi-region sequencing for cancer evolution studies. As the field matures, we expect the method presented here will be incorporated into more analytically comprehensive modelling of whole genome sequencing data when multiple samples are used to infer properties of clonal dynamics. We suggest the next steps are a unified, iterative algorithm that alternates between identifying phylogenetic structure of the constituent clones comprising each tumour sample, and detection of mutations leveraging the new phylogenetic structure. As sequencing costs continue to decrease (e.g. with Illumina’s NovoSeq platform), multi-sample whole genome sequencing of tumours will continue to proliferate as a viable experimental design. Thus, MuClone’s model will be an asset in the arsenal of analytical methods deployed to interpret evolutionary properties of cancer and to gain insights into clonal dynamics in time and space.

## Acknowledgements

This work is supported by long-term funding support from the British Columbia Cancer Foundation. SJ is supported by an NSERC scholarship. SPS and ABC received funding from a Natural Sciences and Engineering Research Council of Canada Frontiers grant #448167-13. SPS received a Canadian Institutes for Health Research Foundation award #FDN 143246 to support this work. SPS holds a Canada Research Chair and is a Michael Smith Foundation for Health Research scholar.

## Supporting Information

**Figure S1.**
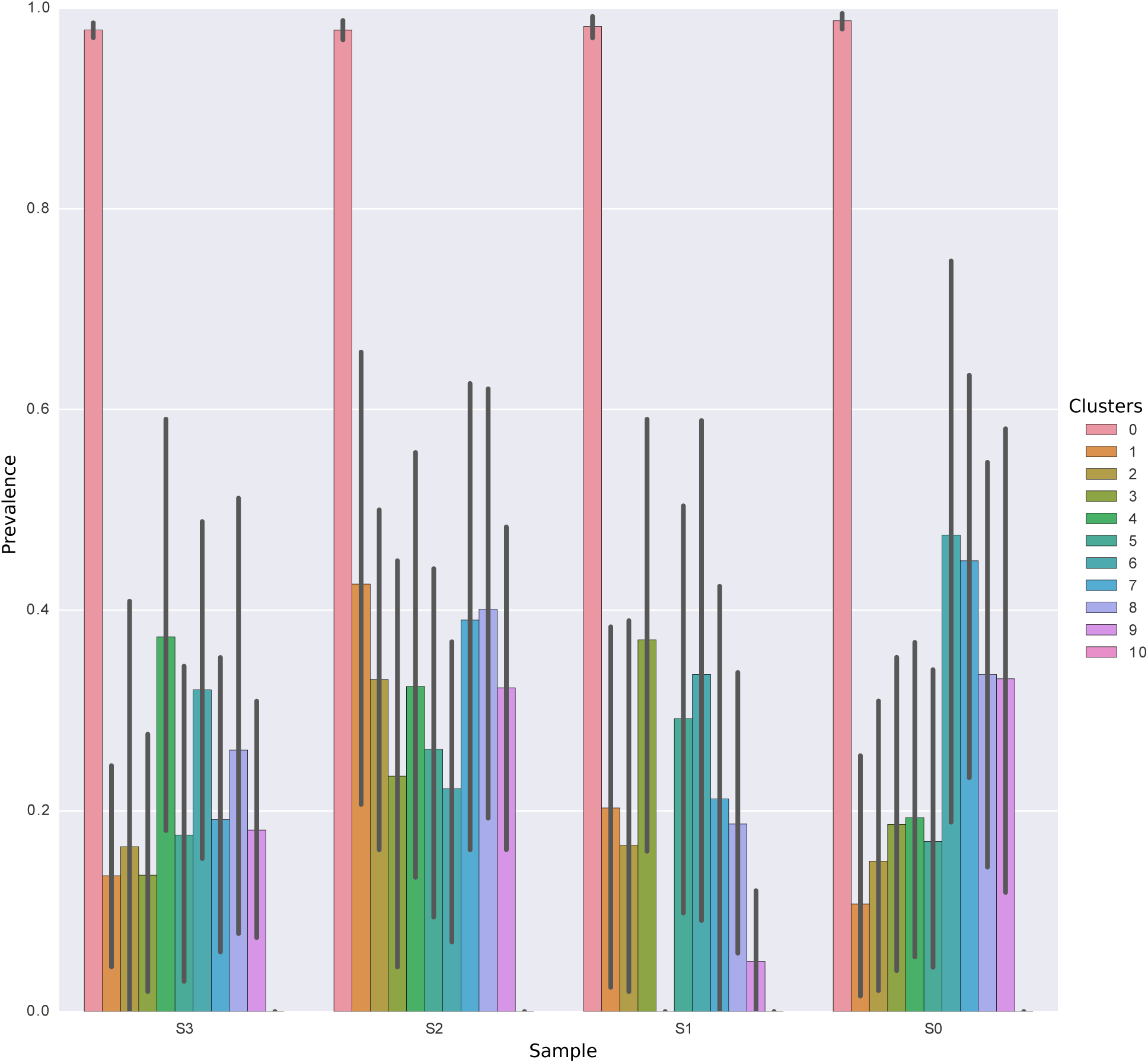
Distribution of the cellular prevalence of clusters across different samples in multiple runs.

**Figure S2.**
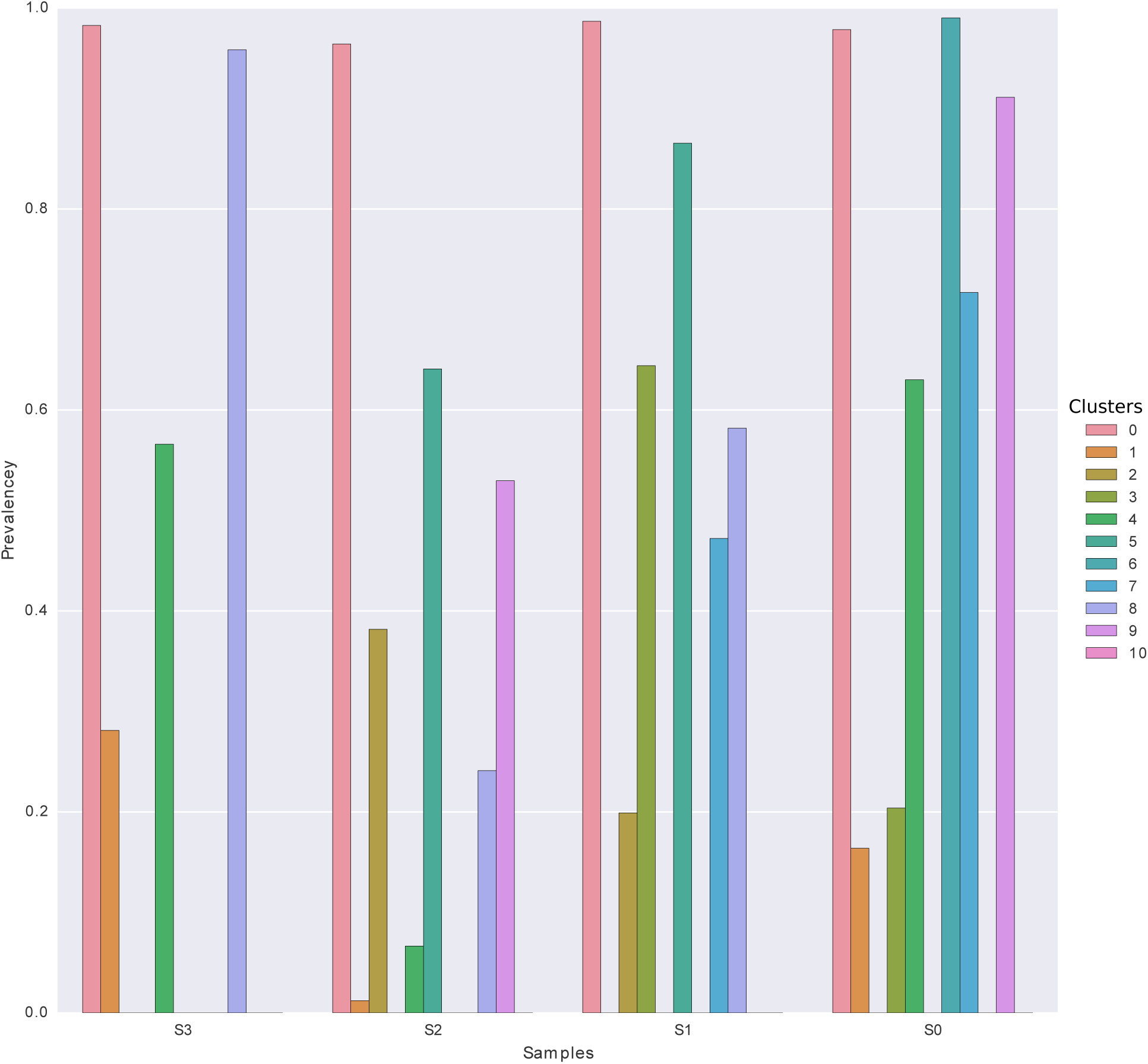
Distribution of the cellular prevalence of clusters across different samples in one random run.

**Figure S3.**
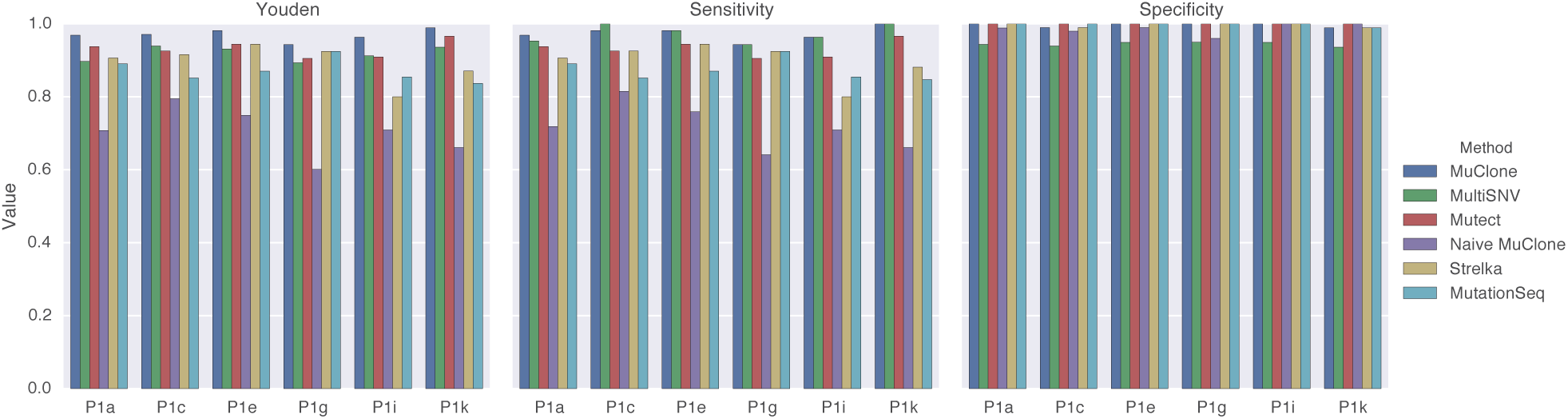
The comparison of Youden’s index, sensitivity and specificity of different mutation detection methods for patient 1. MuClone’s parameters: Wildtype prior is 0.5, ϕ_*T*_ is 0.02, error rate is 0.01, and precision parameter equals 1000.

**Figure S4.**
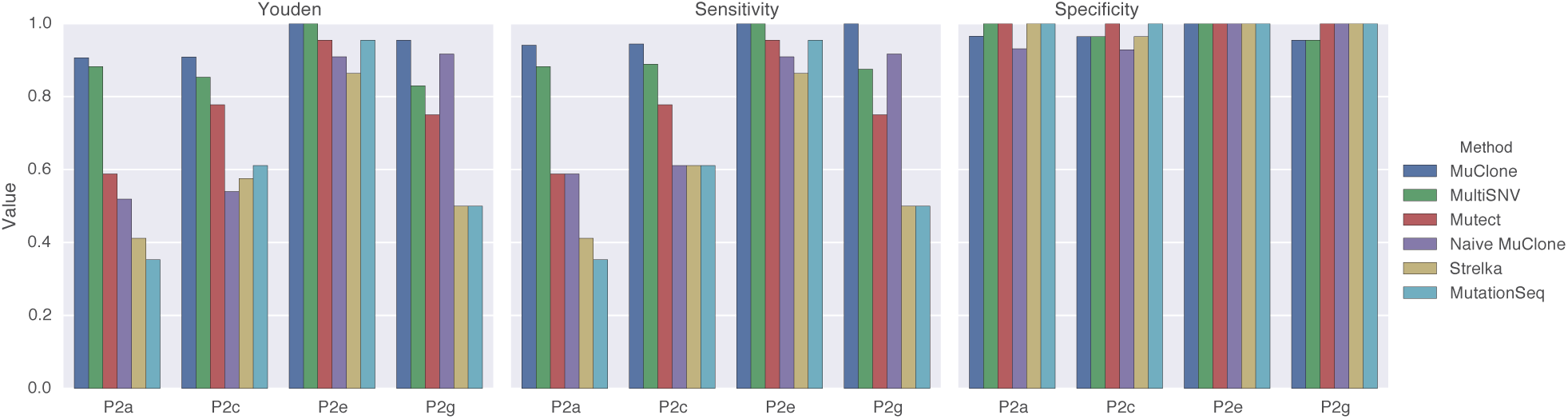
The comparison of Youden’s index, sensitivity and specificity of different mutation detection methods for patient 2. MuClone’s parameters: Wildtype prior is 0.5, ϕ _*T*_ is 0.02, error rate is 0.01, and precision parameter equals 1000.

**Figure S5.**
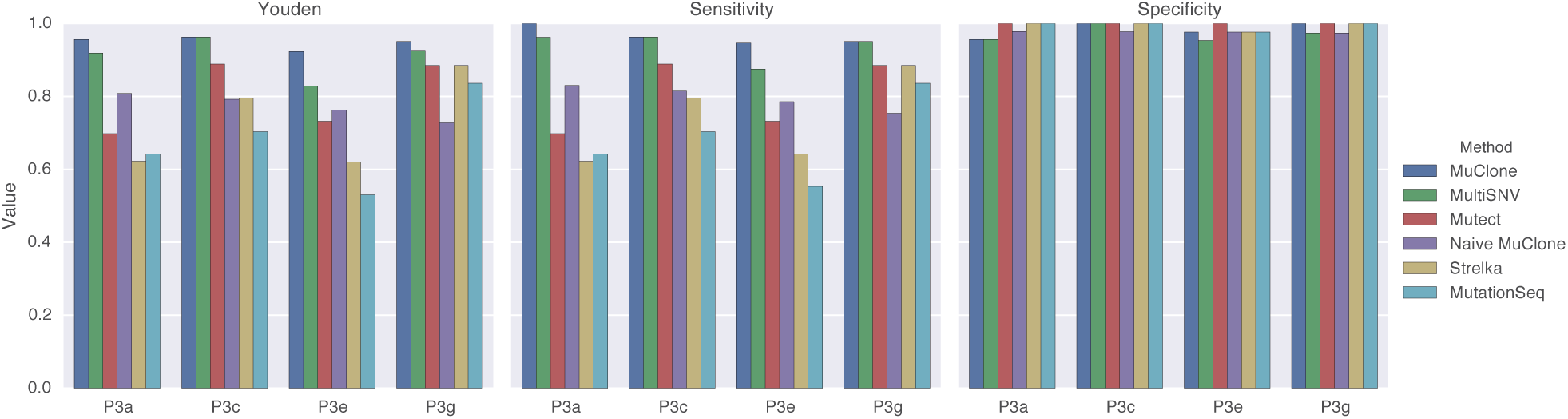
The comparison of Youden’s index, sensitivity and specificity of different mutation detection methods for patient 3. MuClone’s parameters: Wildtype prior is 0.5, ϕ_*T*_ is 0.02, error rate is 0.01, and precision parameter equals 1000.

**Figure S6.**
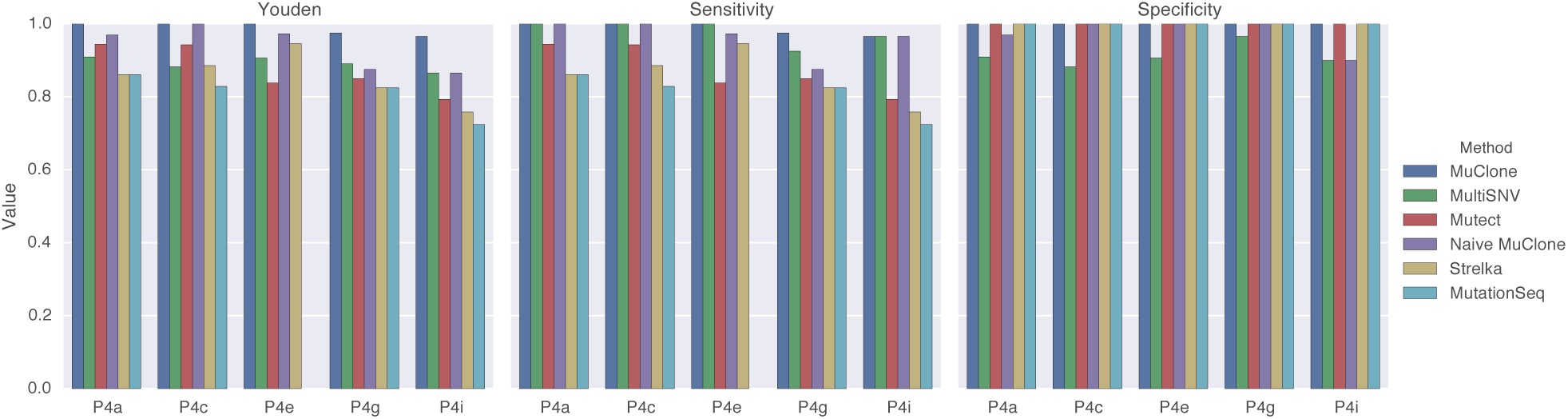
The comparison of Youden’s index, sensitivity and specificity of different mutation detection methods for patient 4. MuClone’s parameters: Wildtype prior is 0.5, ϕ_*T*_ is 0.02, error rate is 0.01, and precision parameter equals 1000.

**Figure S7.**
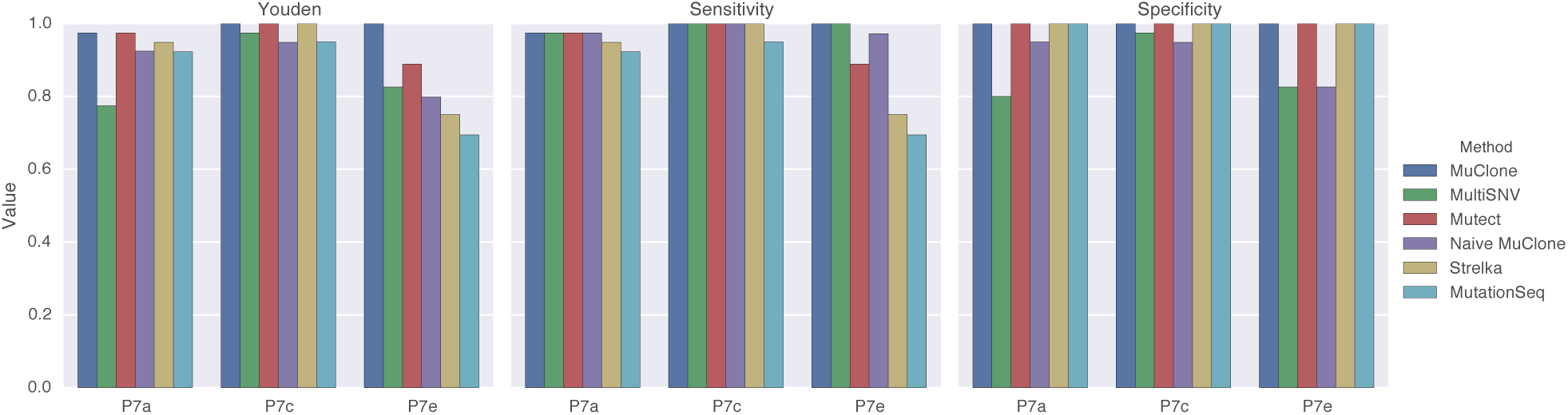
The comparison of Youden’s index, sensitivity and specificity of different mutation detection methods for patient 7. MuClone’s parameters: Wildtype prior is 0.5, ϕ _*T*_ is 0.02, error rate is 0.01, and precision parameter equals 1000.

**Figure S8.**
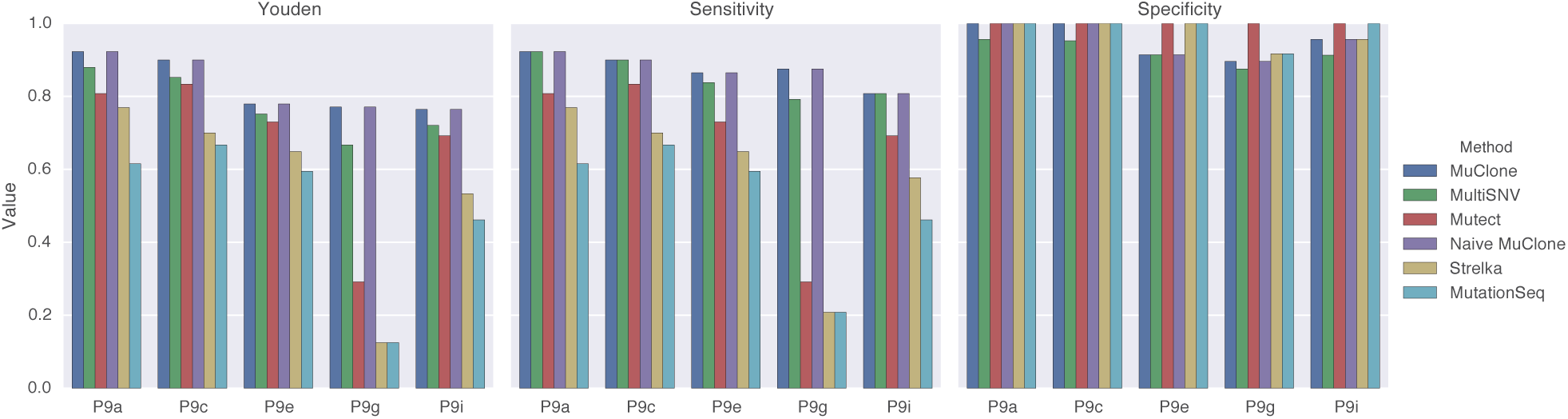
The comparison of Youden’s index, sensitivity and specificity of different mutation detection methods for patient 9. MuClone’s parameters: Wildtype prior is 0.5, ϕ _*T*_ is 0.02, error rate is 0.01, and precision parameter equals 1000.

**Figure S9.**
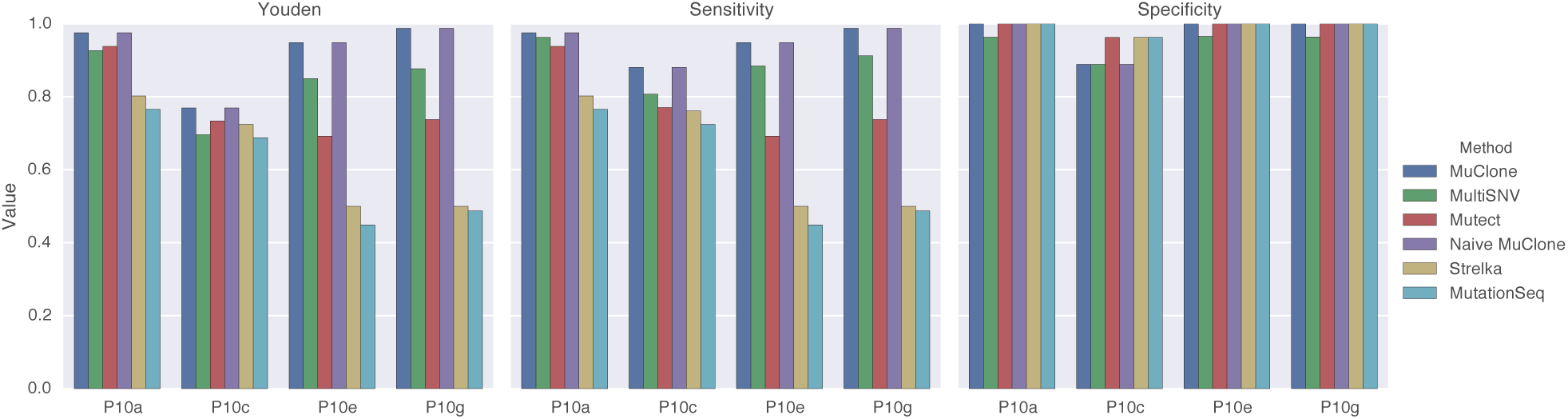
The comparison of Youden’s index, sensitivity and specificity of different mutation detection methods for patient 10. MuClone’s parameters: Wildtype prior is 0.5, ϕ _*T*_ is 0.02, error rate is 0.01, and precision parameter equals 1000.

**Figure S10.**
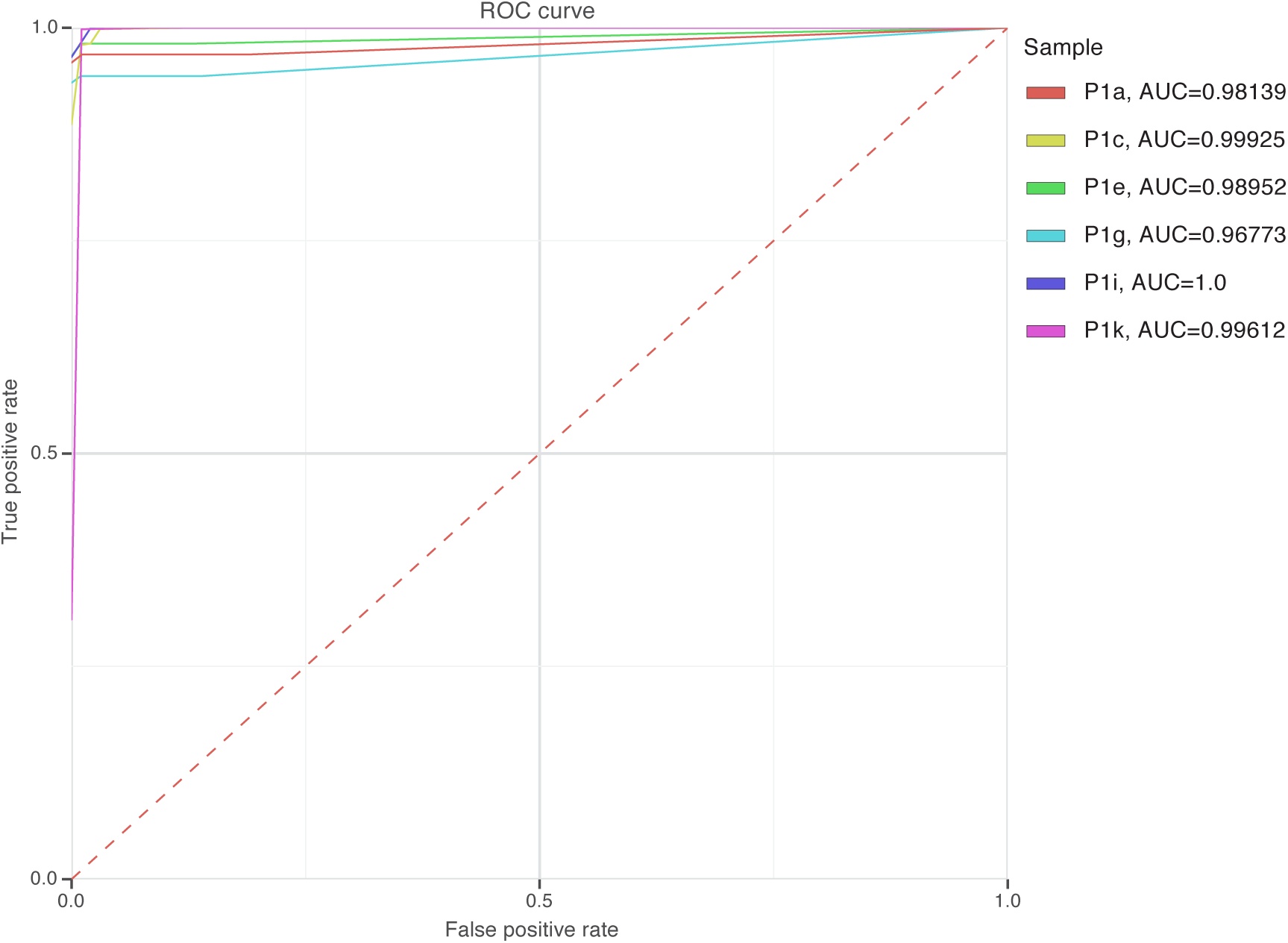
MuClone’s Roc curves and the area under the curve (AUC) for patient 1. MuClone’s parameters: Wildtype prior is 0.5, ϕ _*T*_ is 0.02, error rate is 0.01, and precision parameter equals 1000.

**Figure S11.**
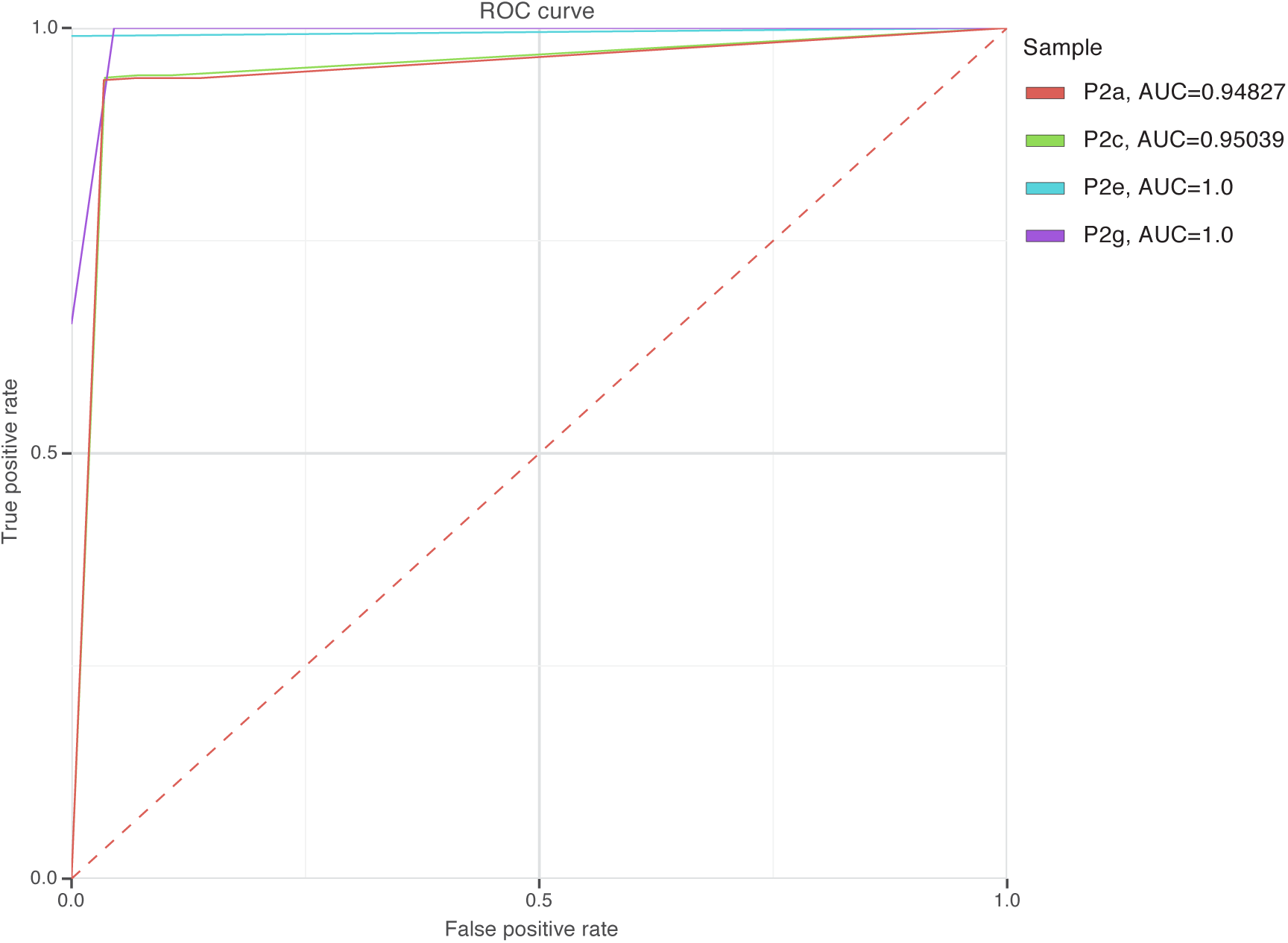
MuClone’s Roc curves and the area under the curve (AUC) for patient 2. MuClone’s parameters: Wildtype prior is 0.5, ϕ _*T*_ is 0.02, error rate is 0.01, and precision parameter equals 1000.

**Figure S12.**
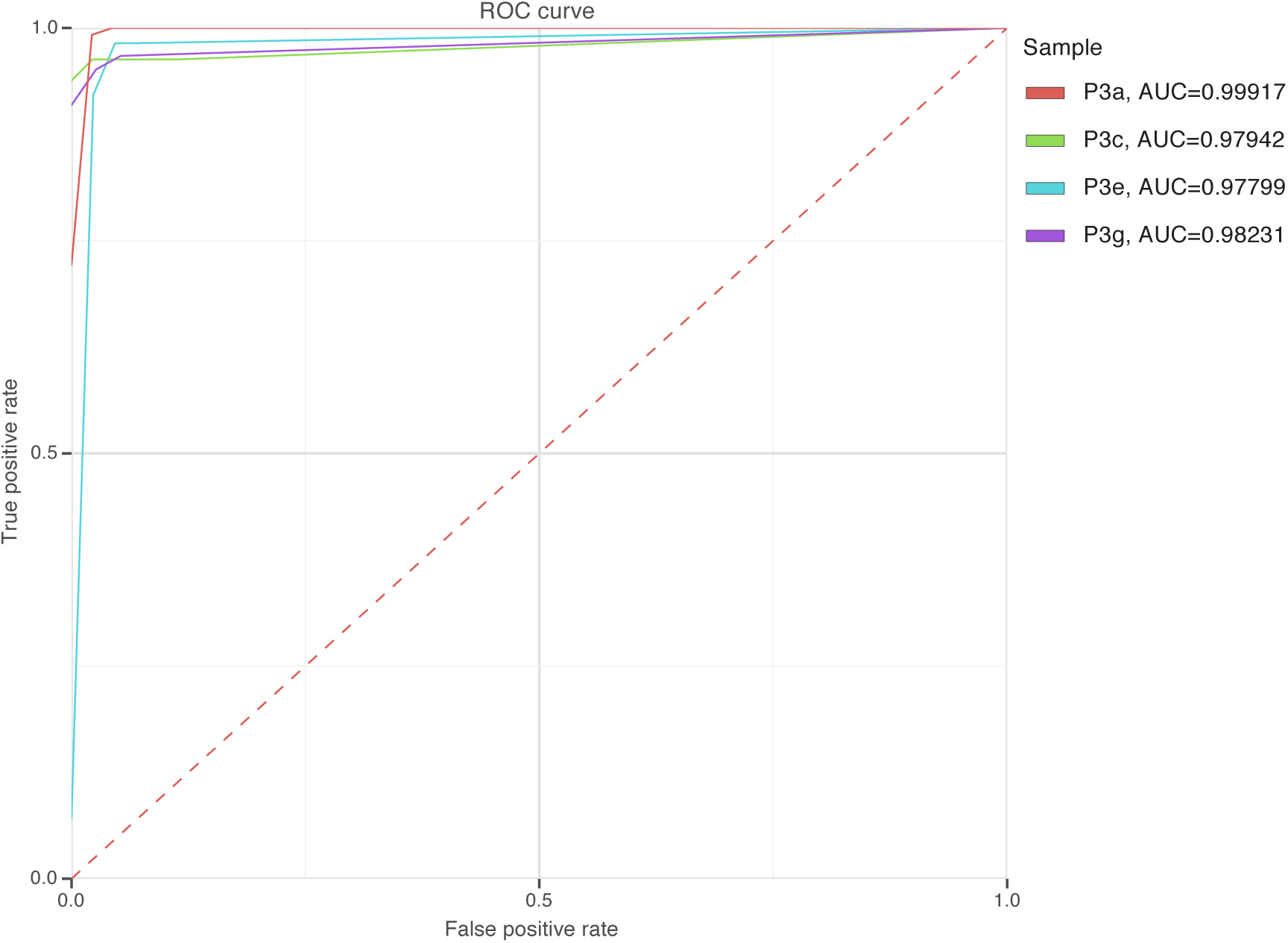
MuClone’s Roc curves and the area under the curve (AUC) for patient 3. MuClone’s parameters: Wildtype prior is 0.5, ϕ _*T*_ is 0.02, error rate is 0.01, and precision parameter equals 1000.

**Figure S13.**
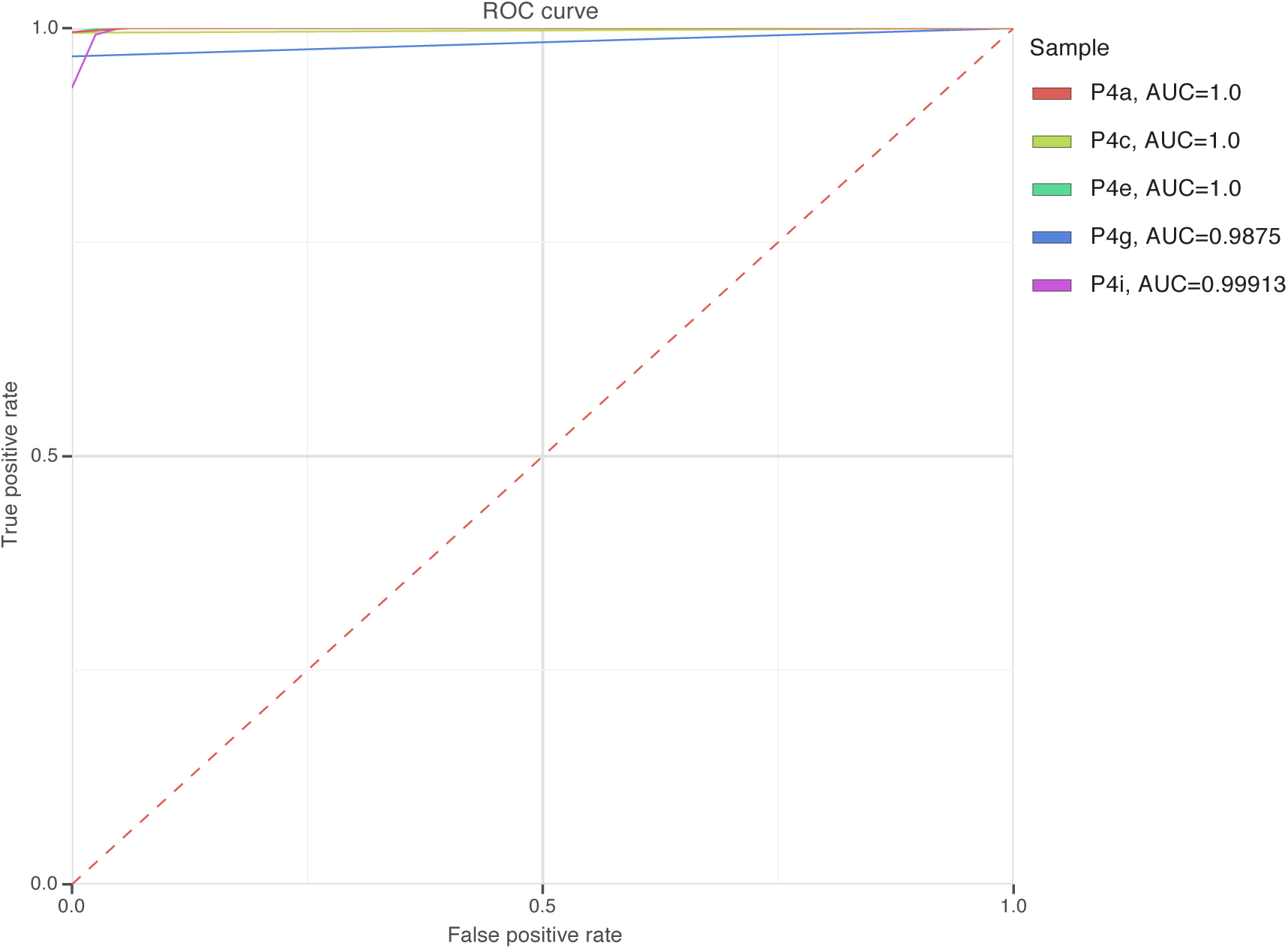
MuClone’s Roc curves and the area under the curve (AUC) for patient 4. MuClone’s parameters: Wildtype prior is 0.5, ϕ _*T*_ is 0.02, error rate is 0.01, and precision parameter equals 1000.

**Figure S14.**
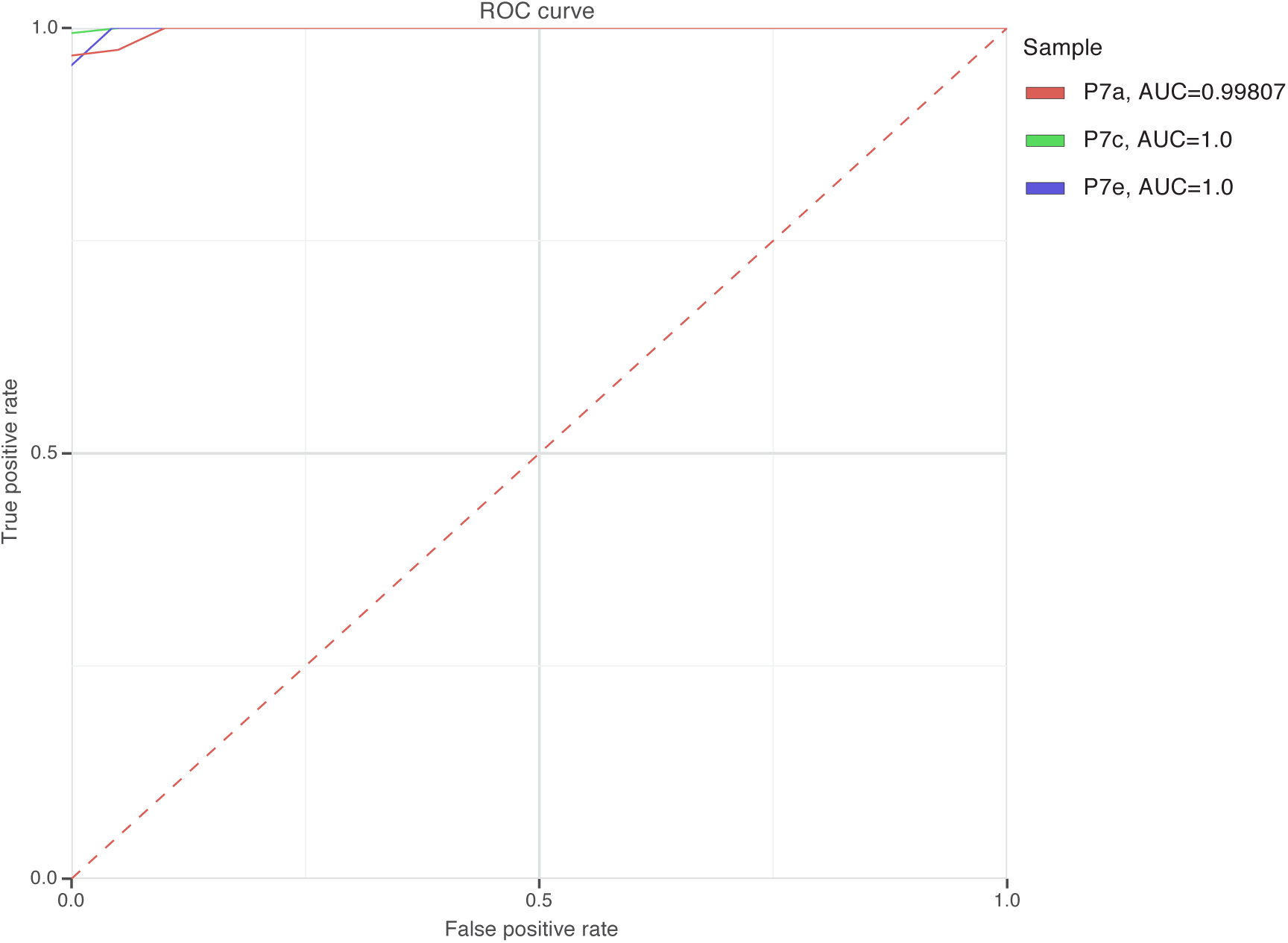
MuClone’s Roc curves and the area under the curve (AUC) for patient 7. MuClone’s parameters: Wildtype prior is 0.5, ϕ _*T*_ is 0.02, error rate is 0.01, and precision parameter equals 1000.

**Figure S15.**
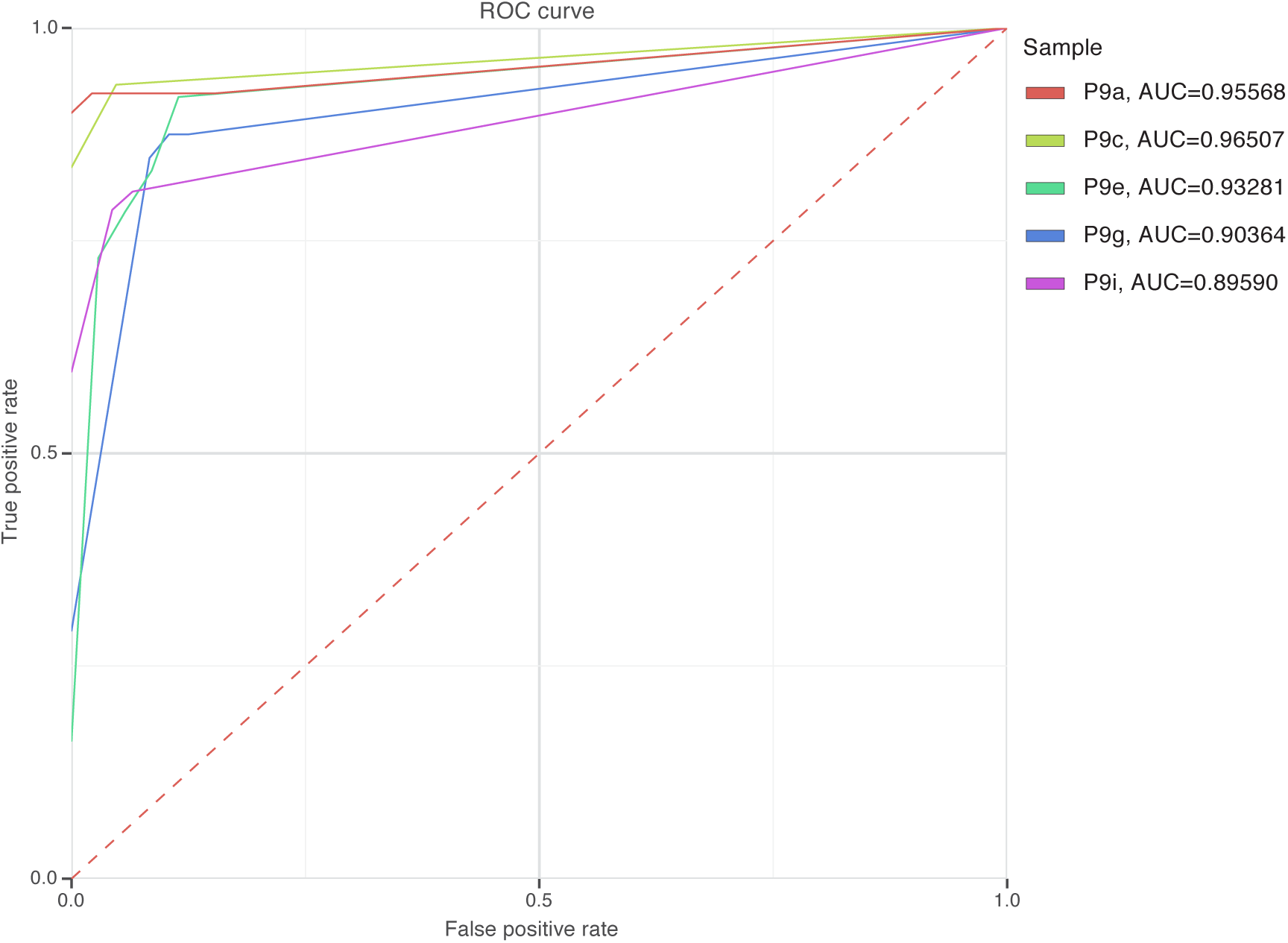
MuClone’s Roc curves and the area under the curve (AUC) for patient 9. MuClone’s parameters: Wildtype prior is 0.5, ϕ _*T*_ is 0.02, error rate is 0.01, and precision parameter equals 1000.

**Figure S16.**
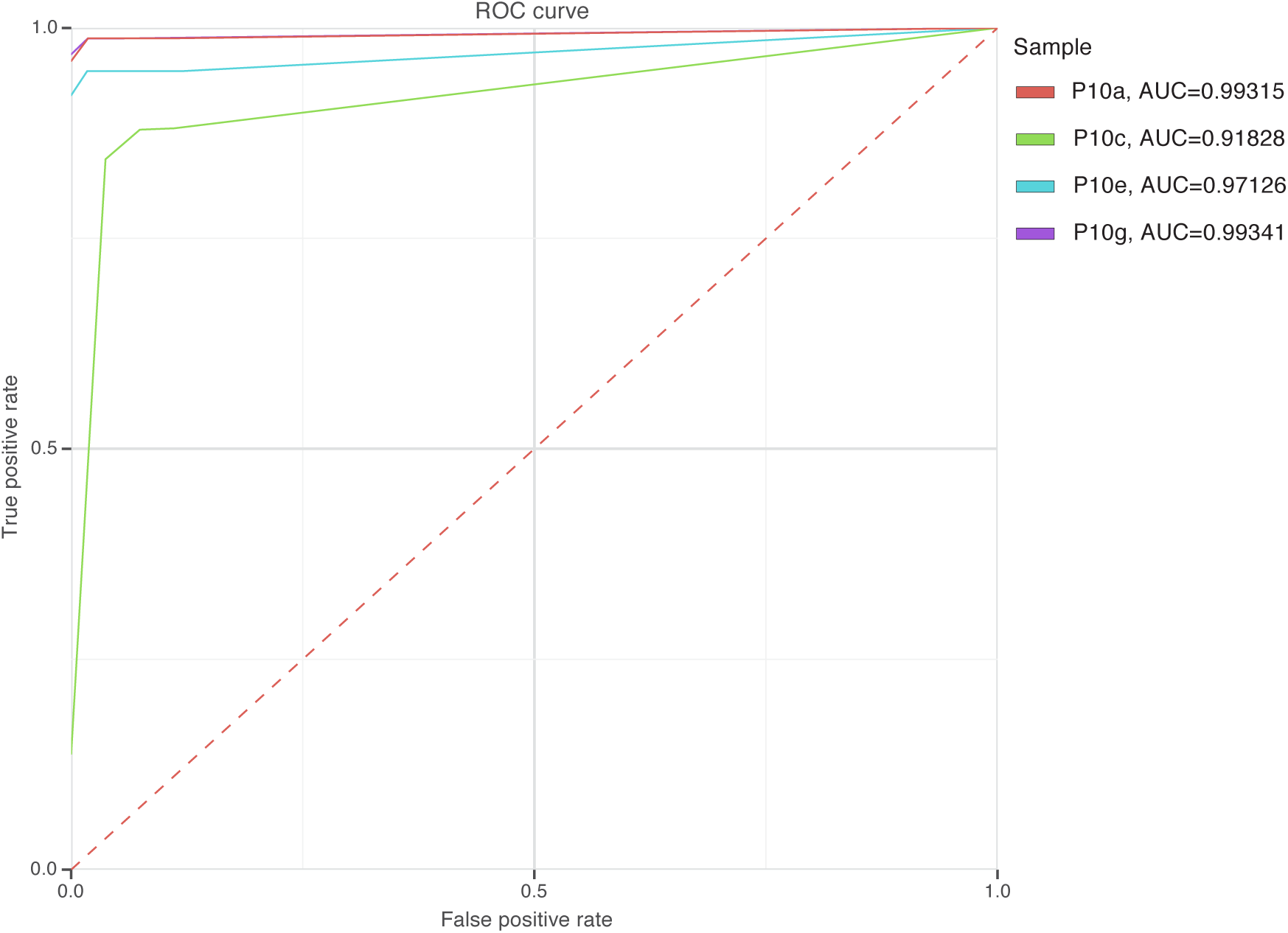
MuClone’s Roc curves and the area under the curve (AUC) for patient 10. MuClone’s parameters: Wildtype prior is 0.5, ϕ _*T*_ is 0.02, error rate is 0.01, and precision parameter equals 1000.

## References

1. K. Cibulskis et al. Sensitive detection of somatic point mutations in impure and heterogeneous cancer samples. Nature Biotechnology, 31(3):213–219, Mar 2013.

2. E. C. De Bruin et al. Spatial and temporal diversity in genomic instability processes defines lung cancer evolution. Science, 346(6206):251–256, 2014.

3. A. G. Deshwar et al. PhyloWGS: Reconstructing subclonal composition and evolution from whole-genome sequencing of tumors. Genome Biology, 16(1):1–20, 2015.

4. J. Ding et al. Feature-based classifiers for somatic mutation detection in tumour-normal paired sequencing data. Bioinformatics, 28(2):167–175, Jan 2012.

5. P. Eirew et al. Dynamics of genomic clones in breast cancer patient xenografts at single-cell resolution. Nature, 518(7539):422–426, 02 2015.

6. M. Gerlinger et al. Intratumor heterogeneity and branched evolution revealed by multiregion sequencing. New England Journal of Medicine, 366(10):883–892, 2012.

7. R. Govindan et al. Genomic landscape of non-small cell lung cancer in smokers and never-smokers. Cell, 150(6):1121–1134, Sep 2012.

8. R. Goya et al. SNVMix: predicting single nucleotide variants from next-generation sequencing of tumors. Bioinformatics, 26(6):730–736, Mar 2010.

9. G. Ha et al. TITAN: inference of copy number architectures in clonal cell populations from tumor whole-genome sequence data. Genome Research, 24(11):1881–1893, 11 2014.

10. M. Jamal-Hanjani et al. Tracking the evolution of non–small-cell lung cancer. New England Journal of Medicine, 367(22):2109–2121, 2017.

11. M. Josephidou et al. MultiSNV: a probabilistic approach for improving detection of somatic point mutations from multiple related tumour samples. Nucleic Acids Research, 43(9):e61, May 2015.

12. S. Kim et al. Virmid: accurate detection of somatic mutations with sample impurity inference. Genome biology, 14(8):R90, 2013.

13. D. C. Koboldt et al. VarScan: variant detection in massively parallel sequencing of individual and pooled samples. Bioinformatics, 25(17):2283–2285, Sep 2009.

14. R. Kridel et al. Histological transformation and progression in follicular lymphoma: A clonal evolution study. PLoS Med., 13(12):e1002197, Dec 2016.

15. A. McKenna et al. The genome analysis toolkit: a map reduce framework for analyzing next-generation DNA sequencing data. Genome Res, 20(9):1297–1303, Sep 2010.

16. A. McPherson et al. Divergent modes of clonal spread and intraperitoneal mixing in high-grade serous ovarian cancer. Nature Genetics, (48):758–767, 2016.

17. S. Nik-Zainal et al. The life history of 21 breast cancers. Cell, 149(5):994–1007, May 2012.

18. V. Popic et al. Fast and scalable inference of multi-sample cancer lineages. Genome Biology, 16(1):1–17, 2015.

19. A. Roth et al. JointSNVMix: a probabilistic model for accurate detection of somatic mutations in normal‵tumour paired next-generation sequencing data. Bioinformatics, 28(7):907–913, Apr 2012.

20. A. Roth et al. PyClone: statistical inference of clonal population structure in cancer. Nature Methods, 11(4):396–398, Apr 2014.

21. R. Salari et al. Inference of tumor phylogenies with improved somatic mutation discovery. Journal of Computational Biology, 20(11):933–944, March 2013.

22. C. T. Saunders et al. Strelka: accurate somatic small-variant calling from sequenced tumor-normal sample pairs. Bioinformatics, 28(14):1811–1817, Jul 2012.

23. S. P. Shah et al. Mutational evolution in a lobular breast tumour profiled at single nucleotide resolution. Nature, 461(7265):809–813, Oct 2009.

24. K. E. Van Rens et al. SNV-PPILP: refined SNV calling for tumor data using perfect phylogenies and ILP. Bioinformatics, 31(7):1133–1135, Apr 2015.

25. S. Yachida et al. Distant metastasis occurs late during the genetic evolution of pancreatic cancer. Nature, 467(7319):1114–1117, Oct 2010.

26. K. Yuan et al. BitPhylogeny: a probabilistic framework for reconstructing intra-tumor phylogenies. Genome Biology, 16:36, 2015.

